# Single-Cell Analysis of NK Cell Cytotoxicity in Cancer Therapy Using Microfluidic Droplets

**DOI:** 10.1101/2025.05.04.651923

**Authors:** Rana S Ozcan, Fatemeh Vahedi, Shina Namakian, Ali A Ashkar, Tohid F Didar

**Affiliations:** School of Biomedical Engineering, McMaster University, Hamilton, Ontario, Canada; McMaster Immunology Research Center, Department of Medicine, McMaster University, Hamilton, Ontario, Canada; Centre for Discovery in Cancer Research, McMaster University, Hamilton, Ontario, Canada

**Keywords:** droplet microfluidics, NK cells, K562, cancer immunotherapy, cytotoxicity testing

## Abstract

Natural Killer (NK) cells are critical components of the immune system, uniquely capable of detecting and eliminating cancer cells without prior sensitization. Here, a droplet-based microfluidic platform is introduced that enables real-time monitoring and single-cell analysis of NK cell-mediated cytotoxicity against K562 cancer cells. Distinct NK cells are evaluated to quantify key metrics, including the percentage of cytotoxic NK cells, serial killing capacity, killing time per target, NK-target contact duration, and migration velocities. The results demonstrated that expanded NK cells (exNK) exhibited longer attachments, superior cytotoxic activity, serial killing, and rapid killing dynamics, whereas peripheral blood NK cells (pbNK), especially when they were exposed to ascites tumor microenvironment (TME) (pbNK-asc), displayed reduced cytotoxic abilities in all parameters. Interestingly, expanded NK cells exposed to ascites TME (exNK-asc) retained partial functionality, indicating that expansion provides resilience against suppressive factors. In addition, cell velocity analysis further revealed that the presence of a cancer cell increases the migration of NK cells. This single-cell analysis provides novel insights into NK-cancer cell interactions, offering a robust framework for enhancing the efficacy of future immunotherapy applications especially for optimizing off-the-shelf NK cell-based immunotherapies.

**Table of Contents:** A droplet-based microfluidic platform examines how Natural Killer (NK) cells target cancer cells at the single-cell level. By comparing peripheral blood and expanded NK cells under normal and tumor-like conditions, distinct differences in attachment, serial killing, killing time, and migration are revealed. These findings provide insights that could enhance tumor targeting, particularly in off-the-shelf NK cell therapies.

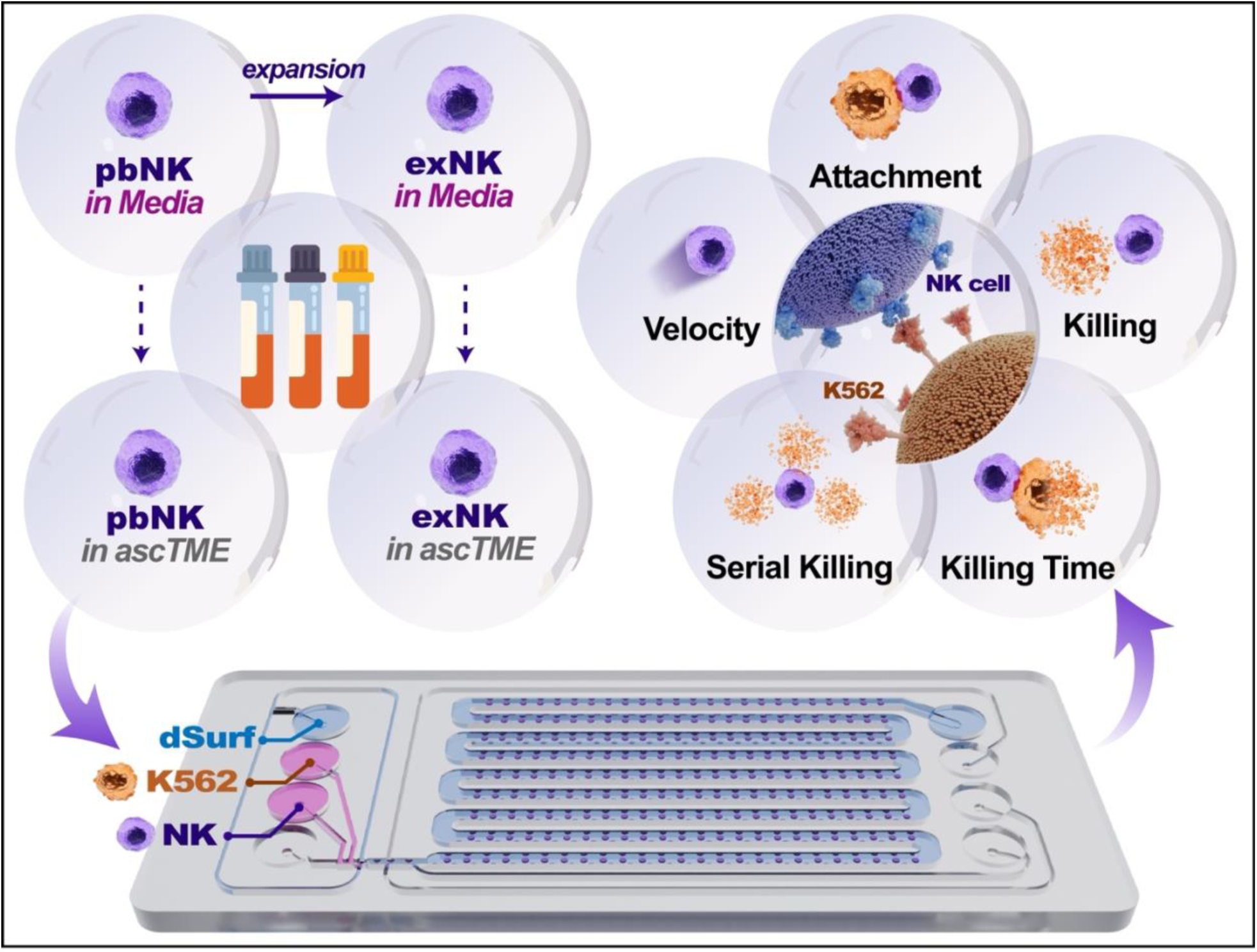

## 1. Introduction

Natural Killer (NK) cells are a key part of the immune system. They are known for their innate cytotoxic capabilities, allowing them to identify and destroy abnormal cells, such as cancerous or virus-infected cells, without prior sensitization while leaving normal cells intact ^1^. Their ability to kill target cells directly and in an antigen independent manner makes them a promising agent in the development of safe off-the-self cancer immunotherapies. Despite their potential, the effectiveness of NK cells can vary based on conditioning, phenotype, and exposure to external stimuli ^2,3^. While NK cells hold promise for a safer and efficient cancer treatment option, traditional methods for monitoring their cytotoxic activity have several limitations. Conventional bulk assays lack the ability to provide single-cell analysis and often fail to capture the nuances of how differently conditioned NK cell phenotypes behave. These limitations create a gap in our understanding, making it challenging to accurately identify the most effective NK cell subpopulations for therapeutic use and optimize their application in immunotherapy ^4^. There is a need for a more precise method that enables the comparison of NK cells in terms of their cytotoxic abilities. To address these gaps, a system that allows the detailed observation and comparison of individual NK cell behaviors under different conditions is essential. Such a system would enable monitoring of NK cell cytotoxicity, including their killing activity, killing time, and other dynamic interactions with cancer cells with real-time monitoring.

Microfluidics devices have significantly advanced cellular research by enabling precise control of the cellular microenvironment ^5–8^, facilitating high-throughput cytotoxicity assays, generation of stable chemical gradients ^9^, single-cell analyses ^10^, efficient cell sorting ^11–13^ and detection. Several droplet-based microfluidic models have been developed in recent years to explore NK cell-mediated cytotoxicity ^14–22^. Existing single-cell investigations show functional heterogeneity among NK cell populations; however, comprehensive comparisons across various conditioning environments are rarely addressed. Many do not thoroughly investigate the effect of varying effector-to-target (E:T) ratios on NK cell efficiency, typically focusing on a narrow range of conditions. Additionally, a detailed examination of diverse NK cell types—such as primary NK cells, expanded NK cells, and those conditioned in tumor microenvironment (TME), such as malignancy ascites, is often missing. Critical aspects like the frequency of NK cell attachments to target cells, their killing efficiency in terms of the number of target cells killed and killing times, and the impact of NK cell velocity on their cytotoxic activity have not been extensively analyzed in these models. Building upon these studies, we investigated NK cell-mediated cytotoxicity against cancer cells using a droplet-based microfluidic platform. We specifically compare different NK cell types—primary NK cells versus expanded NK cells, including those conditioned in TME, to understand their efficacy and behavior. The platform also explores the influence of increasing target cell numbers by analyzing cytotoxicity in droplets at E:T ratios ranging from 1:1 to 1:4. We hypothesized that droplet-based microfluidics will enhance our understanding of NK cell behavior by providing insights into cytotoxic activity at the single-cell level, and that differently conditioned NK cells will show distinct behaviors in their killing efficiency, killing time, and interaction dynamics with target cells, thereby supporting the development of more effective immunotherapies. Our findings reveal that expanded NK cells demonstrate superior cytotoxicity, with rapid killing times, higher killing efficiency, and a notable ability for serial killing. In contrast, primary NK cells conditioned in TME exhibited significantly reduced functionality, including delayed and lower cytotoxicity and no serial killing. It is crucial to address these inhibitory factors in order to improve the efficacy of NK cell therapies.

Meanwhile, the enhanced cytotoxic activity of exNK cells offers a promising approach for improved cancer treatment strategies and establishes them as prime candidates for off-the-shelf NK cell-based immunotherapies.

## 2. Results

### 2.1 Microfluidic droplet technology enables precise, single-cell observation of NK cell cytotoxic events

To explore NK cell-mediated cytotoxicity at single-cell level, we employed a droplet-based microfluidic platform capable of encapsulating effector NK cells with target K562 cancer cells. These assays utilized freshly isolated pbNK cells, as well as pbNK cells treated under specific conditions, including exposure to ascites fluid or expansion with K562 cells expressing membrane-bound IL-21 (See Methods 5.2) (Figure 1a and Supplementary Figure 2). These treatments recapitulate diverse microenvironmental cues: ascites fluid provides a tumor-like milieu rich in soluble factors, while membrane-bound IL-21 stimulation drives NK cell proliferation and potentially augments effector function. By sourcing these differently conditioned NK cells from the same donors, we minimized donor-to-donor variability and could attribute observed functional differences to the conditioning regimen itself rather than to inherent genetic or phenotypic disparities among individuals.

**Figure 1:**
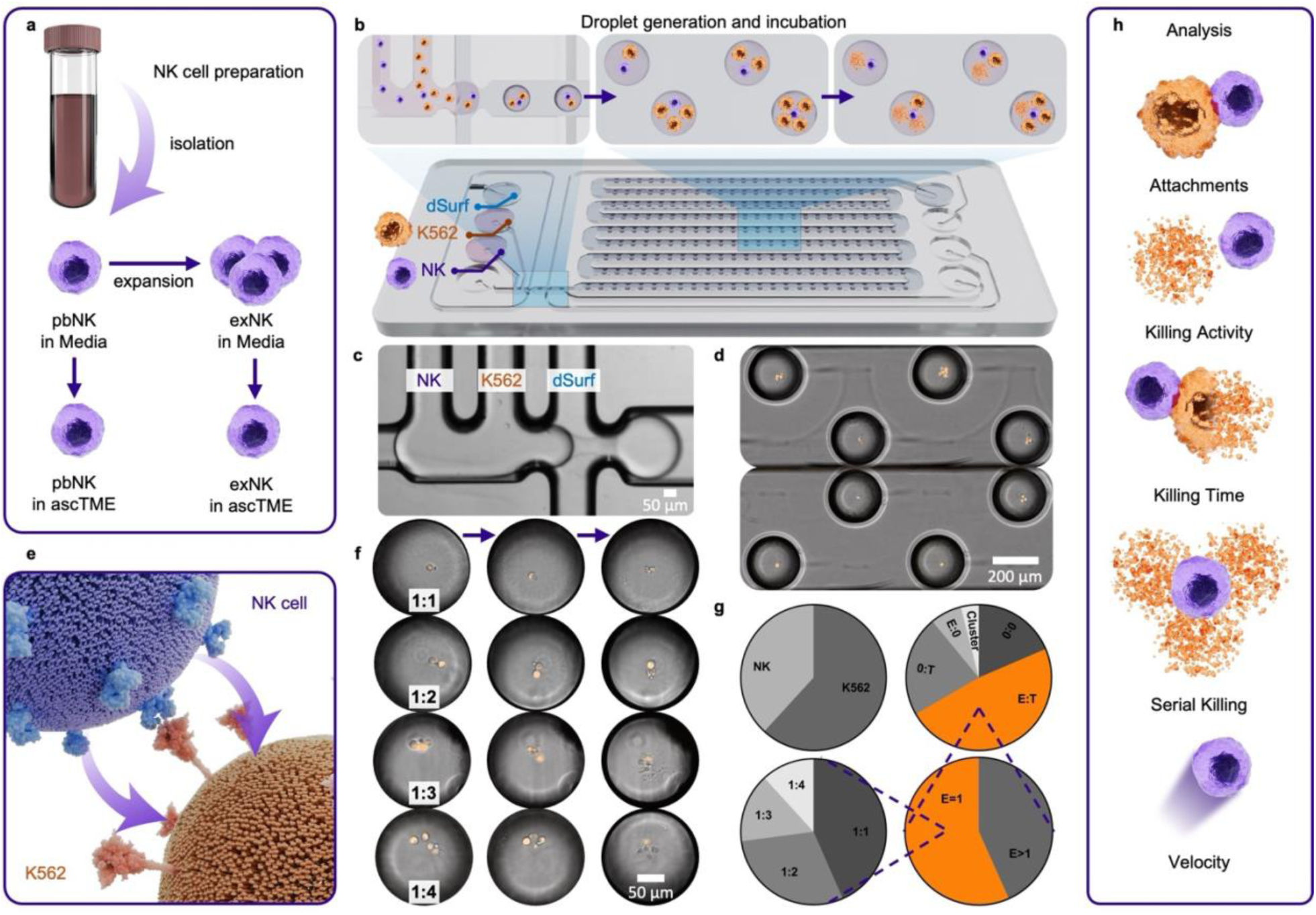
Single-Cell Analysis of NK Cell Cytotoxicity in a Droplet Microfluidic Platform. (a) Schematic illustrating NK cell preparation and (b) subsequent droplet generation and incubation with K562 target cells. (c) Microscopic image showing NK and K562 cells encapsulated in oil-generated droplets and (d) trapped droplets on chip (10X magnification). (e) The interaction of NK and K562 cells. (f) Representative time-lapse images showing NK cell interactions with K562 cells over 10 hours at varying E:T ratios (1:1, 1:2, 1:3, 1:4) (10X magnification). (g) Pie charts depict cell and droplet fractions. (h) Analysis metrics include cell attachments, killing and serial killing activity, killing time, and NK cell velocity.

**Figure 2:**
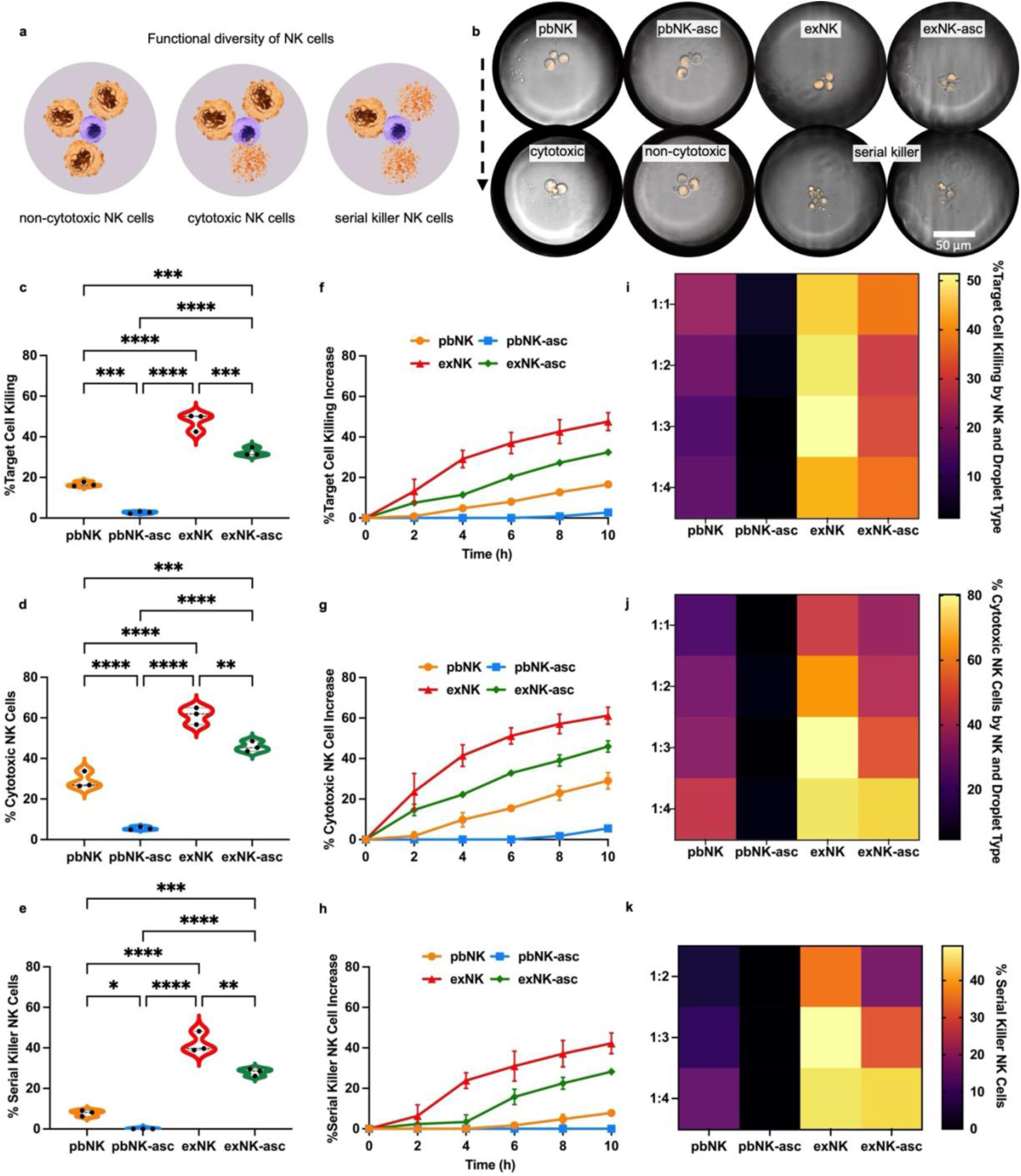
Target cell killing efficiency of NK cells. (a) Schematic showing the functional diversity of NK cells, including non-cytotoxic, cytotoxic, and serial killer NK cells. (b) Representative microscopic images showing NK cell-mediated killing and serial killing activity in droplets across different NK cell types (pbNK, pbNK-asc, exNK, exNK-asc) (10X magnification). Quantification of NK cell activity: (c) percentage of target cell killing, (d) percentage of cytotoxic NK cells, and (e) percentage of serial killer NK cells. Time-dependent cumulative increase in NK cell activity: (f) target cell killing, (g) cytotoxic NK cells, and (h) serial killer NK cells. Heatmaps showing NK cell activity across droplet types: (i) the percentage of target cell killing, (j) the percentage of cytotoxic NK cells, and (k) the percentage of serial killer NK cells at varying E:T ratios. Data represent three independent biological replicates (n=3) for each condition. Asterisks represent “ns” (p > 0.05), * (p ≤ 0.05), ** (p ≤ 0.01), *** (p ≤ 0.001), and **** (p ≤ 0.0001) from one-way ANOVA with Tukey’s multiple comparisons test. All error bars represent the standard deviation.

We utilized the Fluidic 719 (ChipShop) droplet generation and storage chip to perform NK cell-mediated cytotoxicity assays. The chip features a flow-focusing structure with four inlets: one for oil and three for reagents. In our experiments, two inlets were used to introduce K562 target cells and NK effector cells, respectively (Figure 1b and c). Encapsulation efficiencies were optimized to achieve uniform droplet diameters of 175 μm with a volume of 1.5–2 nL. Considering that K562 cells are roughly 20 μm in diameter ^23^ and NK cells are approximately 6 μm ^24^, this droplet size is adequate for our experimental needs. Droplets were generated using input cell concentrations of 2×10^6^ cells/mL for K562 cells and 1×10^6^ cells/mL for NK cells ensuring sufficient space to accommodate the desired cell concentrations for defined E:T ratios ranging from 1:1 to 1:4 (Figure 1c-g). Despite the controlled conditions, variability in cell encapsulation resulted in several droplet categories, including empty droplets, single-cell droplets (only NK or K562), and droplets with multiple NK cells or K562 cells (Figure 1d and g). To address this heterogeneity, after encapsulation, we selected locations with droplets ranging from 1:1 to 1:4 E:T ratios in over 2000 droplet array for analysis. Approximately 48% of droplets contained co-encapsulated NK and K562 cells (E:T droplets) in the chosen locations. Within these E:T droplets 56.7% were classified as desired droplets, containing single NK cells (E=1) interacting with 1–4 K562 cells (Figure 1g). Single-cell analyses are critical to uncovering the intrinsic variability of immune cell function, particularly in the context of cancer immunotherapy. Conventional bulk assays average collective responses, masking the kinetics and cytolytic heterogeneity among individual NK cells. By tracking NK–tumor cell interactions within hundreds of independent droplets, we minimized crosstalk between unrelated cell pairs. Time-lapse microscopy was conducted for 10 hours to monitor these independent NK-K562 interactions and to capture dynamic cytotoxic events at single-cell level (Figure 1e and f). This approach enables real-time observation of critical cytotoxic events, including NK cell attachment to K562 target cells, NK cell-mediated killing and serial killing activity, target cell killing times, and NK cell migration within droplets (Figure 1h).

### 2.2 Droplet microfluidics reveal functional diversity of NK cells through cytotoxicity and serial killing efficiency

The functional diversity of NK cells is highlighted by distinguishing three subsets based on their ability to lyse target cells: non-cytotoxic NK cells, cytotoxic NK cells, and serial killer NK cells (Figure 2a). Non-cytotoxic NK cells fail to eliminate any targets, cytotoxic NK cells kill a single target, and serial killer NK cells are capable of sequentially eliminating multiple targets in a single assay. This stratification is significant for understanding the spectrum of NK cell functionality, where baseline, intermediate, and highly active cells may respond differently in a tumor setting. Visual comparisons of these subsets reveal marked differences in killing behavior: while pbNK-asc droplets generally contain non-cytotoxic cells, exNK droplets frequently show pronounced serial killing events (Figure 2b).

The target cell killing efficiency, defined as the percentage of K562 cells eliminated, revealed clear functional differences across NK cell types (Figure 2c). exNK cells displayed the highest killing capacity, eliminating nearly half of all K562 targets (%47.6), followed closely by exNK-asc with %32.5 killing. In contrast, pbNK showed relatively limited killing, while pbNK-asc exhibited negligible cytotoxicity, %16.6 and %2.7 respectively. These trends were consistent across three biologic replicates. In addition to overall killing, we analyzed cytotoxic NK cells, defined by their ability to eliminate at least one target cell (Figure 2d). The exNK population was the most active, with nearly twice the cytotoxic NK cell frequency compared to pbNK at %61.2 and %29 respectively. exNK-asc retained a notable fraction of active NK cells with %45.8, while the majority of pbNK-asc droplets exhibited no cytotoxic events (%5.5). We next examined serial killing activity, defined as the ability of a single NK cell to kill multiple target cells within the droplet (Figure 2e). Therefore, this data set consists of droplets with 1:1 to 1:4 E:T ratios. exNK again displayed the highest serial killing, with nearly 45% of NK cells. exNK-asc also retained notable serial killing ability, achieving approximately 28%. In comparison, pbNK showed limited serial killing (7.8%), and no serial killing observed in pbNK-asc cells.

Time-course analysis of cumulative target cell killing, and cytotoxic and serial killer NK cell ratios revealed distinct cytotoxic performance among NK cell types (Figure 2f-h). exNK cells displayed rapid increase in all categories within the 2-4 hours when exNK-asc showed a delayed but gradual increase. In contrast, pbNK-asc demonstrated minimal increase in cytotoxicity, and no improvement in serial killing over time. Breaking down the data by E:T ratio provides insight into how NK cells handle increasing target cell numbers under the controlled conditions of the droplet-based assay (Figure 2i–k). The importance of segregating droplets according to E:T ratio lies in capturing the capacity of single NK cells or small NK cell populations to respond to escalating challenge levels. The killing percentage ranking remains consistent across all E:T ratios, with exNK cells consistently demonstrating the highest killing activity Notably, the peak killing activity for exNK cells occurs at an E:T ratio of 1:3, while for other NK cells, the peak is observed at a 1:1 ratio. (Figure 2i and Supplementary Figure 5). This suggests that exNK cells maintain or even enhance their cytotoxic performance as the number of target cells increases, whereas other NK cell groups may become exhausted or less efficient with higher target numbers. Furthermore, the trend of higher cytotoxic and serial killer NK cells percentages in exNK and exNK-asc cells continues across all E:T ratios. (Figure 2g, h and Supplementary Figure 6,7). Notably, the percentages of both cytotoxic and serial killer NK cells slightly increased as the E:T ratio increased in all NK cell groups but not in pbNK-asc cells which remained at less than %5 cytotoxicity and zero serial killing among all E:T ratios. This suggests that only functional NK cells’ cytotoxicity and serial killing events may increase as the target cell number goes higher.

### 2.3 Expanded NK Cells Kill More Rapidly, Whereas TME-Exposed NK Cells Display Slower Cytotoxicity compared to pbNK cells

We conducted experiments analyzing killing times across different NK cell populations to evaluate how NK cell treatment conditions influence the speed and efficiency of target cell killing. Determining the time required for NK cells to kill individual target cells is critical for predicting therapeutic efficacy: rapid killers may exert more immediate control of tumor burden, whereas slower killers might permit tumor escape or adaptation. The comparison of NK cell killing times reveals clear differences based on the NK cell type and treatment conditions. Figure 3a provides a schematic representation of NK cell-mediated killing events at different time points. Microscopic images illustrate the differences in killing times across the four NK cell types. The images show representative snapshots of target cell killing at varying time points (Figure 3b). These microscopic observations align with the quantitative data, visually emphasizing the enhanced killing capacity of expanded NK cells (exNK and exNK-asc) compared to the slower activity of peripheral blood NK cells (pbNK and pbNK-asc), particularly under suppression conditions.

**Figure 3:**
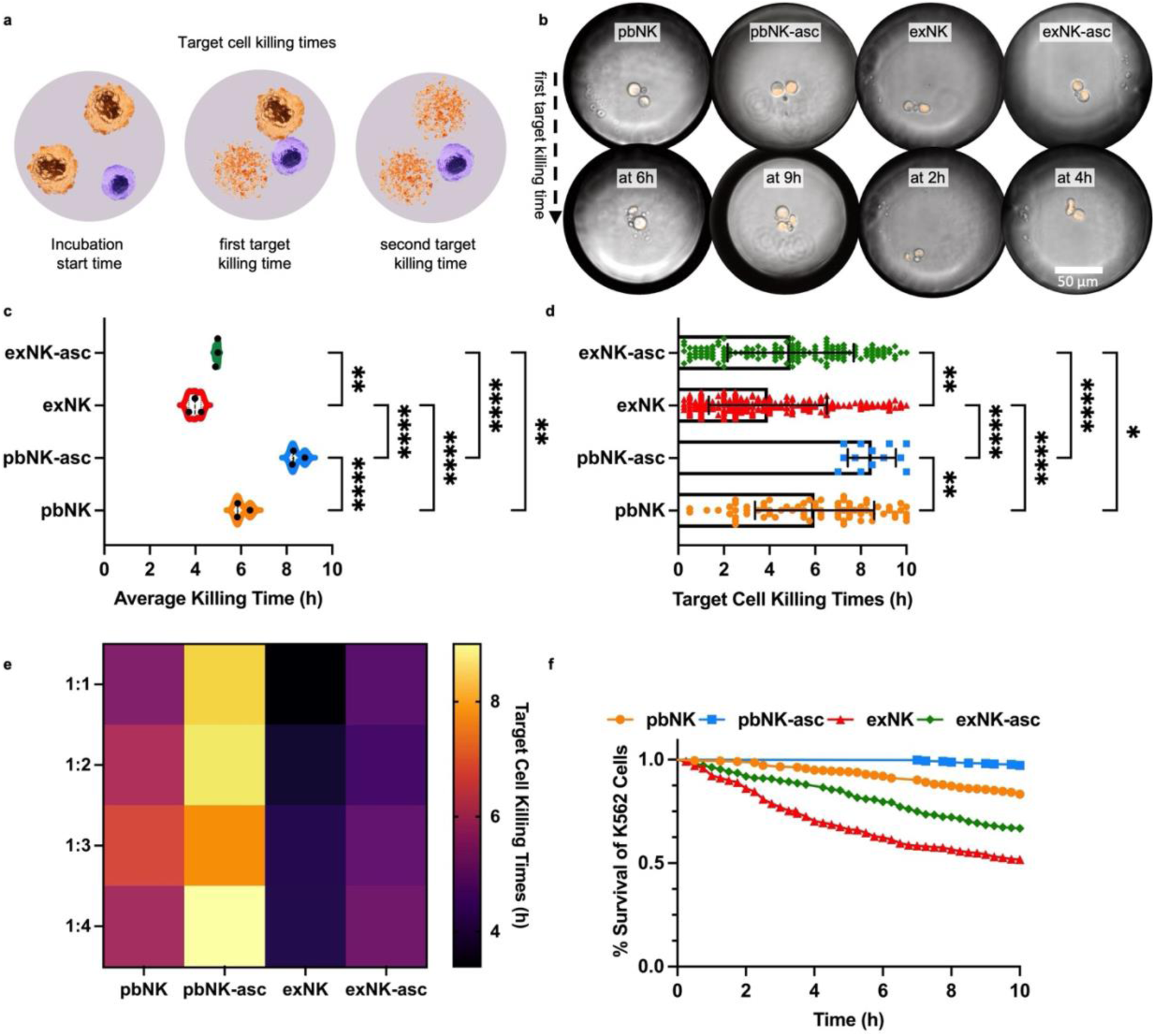
Comparison of NK Cell Killing Times. (a) Illustration of NK cell-mediated killing events at different time points. (b) Representative microscopic images showing NK cell-mediated killing of K562 cells at different time points (10X magnification). (c) Comparison of average target cell killing times among NK cell types. (d) Distribution of target cell killing times across different NK cell types. (e) Heatmap representing target cell killing times across Effector-to-Target (E:T) ratios and NK cell types. (f) K562 cell survival curves over time when co-encapsulated with different NK cell types. Data represent three independent biological replicates (n=3) for each condition. Asterisks represent “ns” (p > 0.05), * (p ≤ 0.05), ** (p ≤ 0.01), *** (p ≤ 0.001), and **** (p ≤ 0.0001) from one-way ANOVA with Tukey’s multiple comparisons test. All error bars represent the standard deviation.

Over a 10-hour incubation period, exNK cells exhibited the fastest average killing time of approximately 4 hours, highlighting their superior cytotoxic efficacy. These cells were followed by exNK-asc cells, which averaged 5 hours to kill K562 target cells. In contrast, pbNK cells showed a slower response, with an average killing time of 6 hours, while pbNK-asc cells demonstrated the longest killing time at 8.5 hours (Figure 3c and Supplementary Figure 9a). This significant delay in pbNK-asc cells indicates that suppression conditions (ascitic fluid) considerably impair their cytotoxic function. The differences between all groups were statistically significant, as indicated by p-values (p ≤ 0.001 or lower), confirming the impact of expansion and suppression treatments on NK cell performance.

The distribution of target cell killing times further emphasizes these trends. exNK cells demonstrated a rapid cytotoxic response, killing most target cells within the first 4 hours (Figure 3d, and Supplementary Figure 9b). This result indicates an efficient and consistent killing capacity following expansion. On the other hand, pbNK-asc cells showed a delayed killing pattern, with a significant portion of target cell death occurring in the 8-10-hour range. This prolonged response aligns with the observed reduction in functionality under suppression conditions, further supporting the impaired killing potential of pbNK-asc cells. The killing times were also analyzed across different E:T ratios to evaluate the impact of target cell density (Figure 3e). At all E:T ratios tested the trends remained consistent: exNK cells consistently demonstrated the fastest killing times, while pbNK-asc cells had the slowest. A slight increase in killing time was observed as the number of target cells increased. This increase is expected, as the presence of more target cells naturally extends the overall killing time.

The survival curves of K562 cells provide further confirmation of these findings (Figure 3f). pbNK-asc cells displayed minimal cytotoxic activity, with the survival percentage remaining close to 100% over the 10-hour period. In contrast, the exNK cells exhibited the most rapid reduction in K562 survival, with nearly 50% of target cells killed over time. pbNK cells and exNK-asc cells displayed intermediate cytotoxicity, following the same trends observed in the average and distributed killing times.

### 2.4 Attachment dynamics reflect functional diversity of NK cells

To investigate how NK cell attachment behavior correlates with their functional diversity, we analyzed attachment dynamics under varying treatment conditions. Attachments are critical for cytotoxicity because an NK cell must establish contact with a tumor cell in order to form an immunological synapse—a specialized interface that enables the directed release of cytotoxic granules and initiates target cell death. The attachment behavior of NK cells toward K562 target cells varied across NK cell types and treatment conditions. Figure 4a illustrates the different outcomes of NK cell interactions with K562 target cells, categorized as no attachment, attachment without killing, and attachment leading to killing. These stages highlight the sequential steps of NK cell-mediated cytotoxicity, starting with the initial recognition and attachment of target cells, followed by the potential for cytotoxic activity. Microscopic images provide representative examples of NK cell behaviors across different conditions. peripheral blood NK cells often showed no attachment and only attachment whereas expanded NK cell groups displayed clear examples of attachment leading to killing, with visible evidence of target cell elimination (Figure 4b).

**Figure 4:**
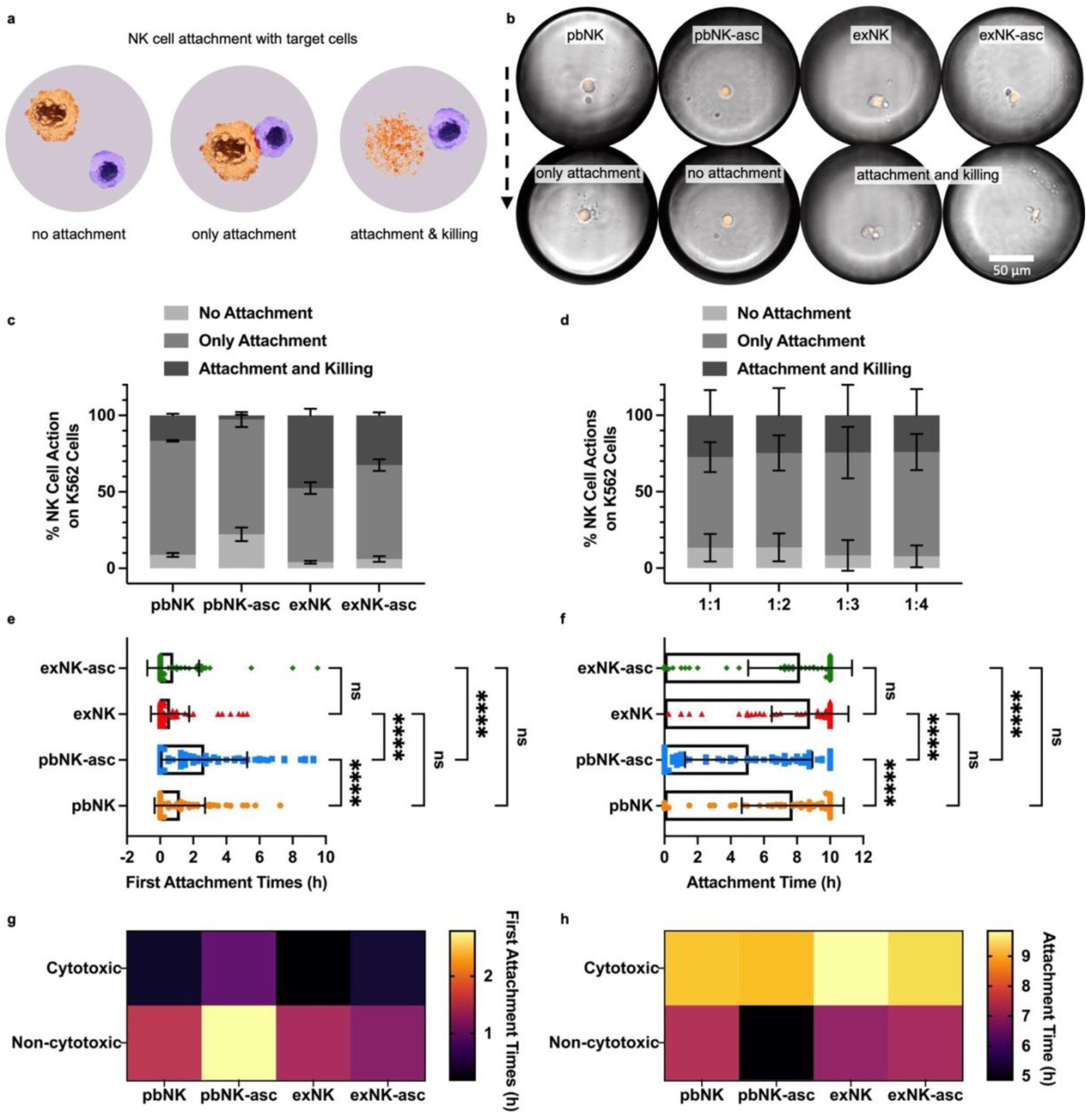
Comparison of NK Cell Attachments with K562 Cells. (a) Schematic illustrating NK cell actions: no attachment, attachment without killing, and attachment leading to killing of K562 cells. (b) Representative microscopic images showing NK cell attachments with K562 cells (10X magnification). Comparison of NK cell actions on K562 target cells (c) based on NK cell type (d) and droplet type. Each bar represents the percentage of NK cells exhibiting no attachment, only attachment, and attachment and killing. (e) First attachment times of NK cells with K562 cells. (f) Time NK cells spent as attached to K562 cells. (g) First attachment times of cytotoxic and non-cytotoxic NK cells with K562 cells across NK cell types. (h) Total time cytotoxic and non-cytotoxic NK cells remained attached to K562 cells across NK cell types. Data represent three independent biological replicates (n=3) for each condition. Asterisks represent “ns” (p > 0.05), * (p ≤ 0.05), ** (p ≤ 0.01), *** (p ≤ 0.001), and **** (p ≤ 0.0001) from one-way ANOVA with Tukey’s multiple comparisons test. All error bars represent the standard deviation.

**Figure 5:**
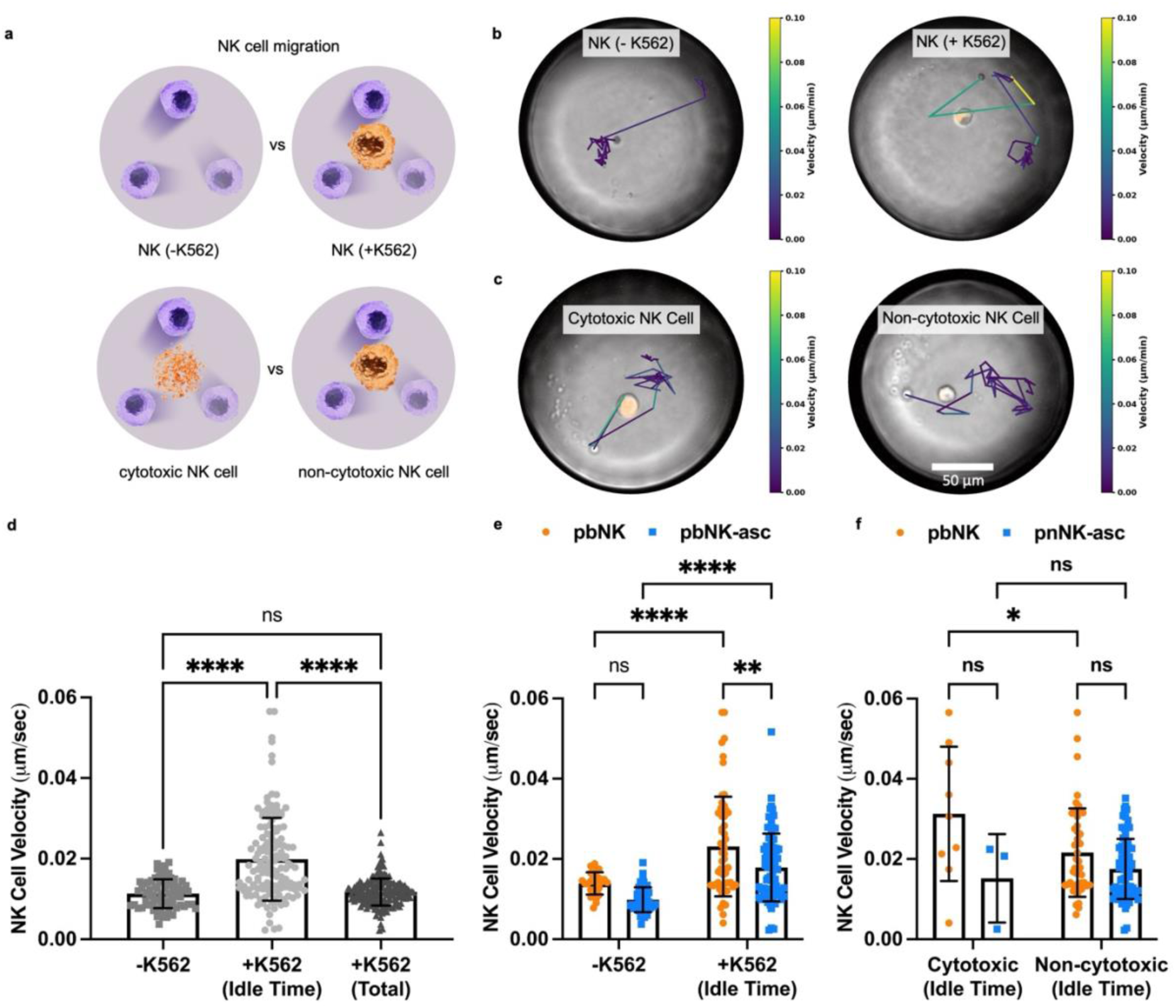
Comparison of NK Cell Migration in Droplets. (a) Schematic depicting NK cell migration under different conditions: with or without K562 cells and comparing cytotoxic versus non-cytotoxic NK cells. (b-c) Representative NK cell migration tracks within droplets. The colormap indicates the instantaneous velocity of the migrating NK cells (10X magnification). (d) NK cell velocity with or without K562 cells, including total and idle times. (e) Comparison of NK cell velocities between pbNK and pbNK-asc under conditions with and without K562 cells. (f) NK cell velocity comparison for cytotoxic and non-cytotoxic NK cells during idle times. Data represent three independent biological replicates (n=3) for each condition. Asterisks represent “ns” (p > 0.05), * (p ≤ 0.05), ** (p ≤ 0.01), *** (p ≤ 0.001), and **** (p ≤ 0.0001) from one-way or two-way ANOVA with Tukey’s multiple comparisons test. All error bars represent the standard deviation.

pbNK-asc cells exhibited the highest percentage of “no attachment” events and the lowest percentage of “attachment and killing”, indicating impaired target recognition and cytotoxic function under suppression conditions. In contrast, exNK cells showed the most effective interaction with target cells, with the lowest “no attachment” percentage and the highest “attachment and killing” percentage (Figure 4c). These results suggest that expansion treatments greatly enhance NK cell attachment efficiency and subsequent cytotoxic activity, while ascites fluid suppresses this capacity. When NK cell actions were compared across different E:T ratios the distribution of NK cell actions remained consistent (Figure 4d). The differences across ratios were minimal, across all tested ratios, indicating that NK cell functionality is maintained regardless of target cell concentration. However, the percentage of “no attachment” events slightly decreased as the number of target cells increased. This decrease is logical, as a higher density of K562 cells increases the likelihood of NK cells locating and attaching to a target cell.

Determining both the first attachment time and the overall duration of attachment provides insights into how quickly NK cells can identify and engage a target cell, as well as how effectively they maintain contact to exert lethal activity. First attachment times of NK cells with K562 target cells are compared in Figure 4e. expanded NK cell groups demonstrated the fastest attachment, with an average first attachment time of less than an hour. This rapid attachment underscores the enhanced target recognition and interaction capacity achieved through expansion. In contrast, pbNK cells required an average of 2 hours to make their first attachment, while pbNK-asc cells exhibited the slowest response, with a significantly delayed first attachment time of 2.7 hours. The first attachment times of pbNK-asc cells were significantly delayed compared to the other groups, while no significant differences were observed among the remaining groups. This indicates that suppression treatments impair the ability of peripheral blood NK cells to locate and attach to target cells quickly. Figure 4f represents the time NK cells spent attached to K562 target cells, revealing additional insights into their functional behavior. exNK cells maintained attachment for the longest duration, averaging 8.8 hours, while exNK-asc cells showed a slightly shorter attachment time of 8.2 hours. In contrast, pbNK cells displayed an average attachment time of 7.7 hours, and ascitic fluid condition had a significant impact on pbNK-asc cells, which exhibited the shortest attachment time at 5 hours.

The first attachment times and attachment durations of cytotoxic and non-cytotoxic NK cells are compared across different NK cell groups (Figure 4g and h). In all groups, cytotoxic NK cells attached significantly earlier than non-cytotoxic NK cells. This highlights the importance of timely attachment for effective target cell killing. Notably, while pbNK, exNK, and exNK-asc cells showed similar trends, pbNK-asc cells displayed a distinct delay, with both cytotoxic and non-cytotoxic subsets attaching later (Figure 4g and Supplementary Figure 10a). Similarly, across all groups, cytotoxic NK cells remained attached to target cells for a significantly longer time compared to non-cytotoxic NK cells. This prolonged attachment likely supports their ability to kill target cells effectively. Furthermore, pbNK, exNK, and exNK-asc cells exhibited comparable trends, whereas pbNK-asc cells showed a significantly shorter attachment time (Figure 4h and Supplementary Figure 10b). Together, these findings demonstrate that while cytotoxic and non-cytotoxic NK cells consistently differ in attachment behavior, pbNK-asc cells uniquely exhibit impaired attachment times and durations which reflects their reduced cytotoxic potential under suppression conditions.

### 2.5 NK cell migration patterns are enhanced in the presence of target cells

Monitoring the velocity of NK cells under varying conditions addresses a key question in immunotherapy research: more motile NK cells may more swiftly locate and engage target cells, leading to faster tumor clearance. The velocity of NK cells under various conditions was analyzed to evaluate their migratory behavior (Figure 5a). Representative heatmaps of NK cell migration within droplets display their instantaneous velocity over a 10-hour period (Figure 5b, c and Supplementary Figure 11). NK cells in the presence of K562 target cells (NK(+K562)) exhibited faster migration compared to NK cells alone (NK(-K562)) until an attachment event occurred. Similarly, cytotoxic NK cells displayed slightly higher velocities compared to non-cytotoxic NK cells prior to target cell attachment (See Supplementary Movie 3). However, once attachment occurred, the NK cells exhibited reduced movement, explaining why the total velocity of NK(+K562) cells was not significantly different from NK(-K562) cells, as shown in Figure 5d.

To account for this reduction, idle time velocities, the velocity of NK cells when no attachment occurred, were analyzed separately (Figure 5e and f). Figure 5e compares the idle time velocities of NK cells across pbNK and pbNK-asc groups. For pbNK cells, the idle time velocity (0.023 μm/sec) was significantly higher than that of NK cells alone (0.014 μm/sec), demonstrating increased motility in the presence of target cells. Similarly, for pbNK-asc cells, the idle time velocity (0.018 μm/sec) was significantly higher compared to the velocity of NK-asc cells alone (0.010 μm/sec). However, the velocity difference between the two types of alone peripheral blood NK cells was not significant.

The idle time velocities of cytotoxic and non-cytotoxic NK cells within the peripheral blood NK cell group were compared in Figure 5f. A significant difference was observed only between cytotoxic and non-cytotoxic pbNK cells, where cytotoxic NK cells exhibited higher velocities. No significant differences were found for pbNK-asc cells. This might suggest that suppression diminishes the velocity advantage of cytotoxic NK cells. Overall, these findings demonstrate that NK cells migrate faster in the presence of target cells until attachment occurs, with cytotoxic NK cells might exhibit higher idle time velocities compared to non-cytotoxic NK cells.

## 3. Discussion

NK cell immunotherapy has emerged as a promising approach in cancer treatment due to the innate cytotoxic capabilities of NK cells, which allow them to identify and destroy cancer cells without prior sensitization ^25^. Accurate assessment of immune cell cytotoxicity is crucial for advancing NK cell-based therapies. Traditional methods, such as the Cr-51 and flow cytometry-based tests, have provided foundational insights but are limited by their inability to offer individual cell analysis and real-time monitoring ^26^. In this study, we utilized droplet-based microfluidic technology to investigate NK cell-mediated cytotoxicity at the single-cell level, providing real-time insights into NK cell interactions with cancer cells, including target cell attachment, killing efficiency, serial killing activity, and migration dynamics. This platform offers significant advantages over traditional cytotoxicity assays by enabling precise, high-throughput, single-cell analysis and addressing critical limitations such as lack of real-time monitoring and inability to capture functional heterogeneity within NK cell populations ^14,20–22^ (Supplementary Movie 2).

The findings from this study provided significant insights into the functionality of NK cells under various conditions and underlined the potential of NK cell-based immunotherapies. exNK cells demonstrated the highest cytotoxic activity, including the fastest killing times (∼4 hours), the highest killing efficiency (47.6%), and notable serial killing activity (∼45%). These findings align with prior studies showing that expansion processes enhance NK cell functionality and anti-tumor activity by improving their cytotoxic potential and proliferative capacity ^27–29^. The ability of exNK cells to maintain or even improve their killing efficiency at higher E:T ratios (peaking at 1:3) This robustness positions them as strong candidates for developing NK cell-based treatments for cancer.

Conversely, pbNK-asc cells exhibited significantly reduced functionality, with minimal cytotoxic activity (2.7%) and no serial killing events observed. These cells showed delayed first attachment times (2.7 hours) and shorter attachment durations (∼5 hours). This reflecs the profound negative effects of suppression conditions, such as exposure to the ascites tumor microenvironment. This aligns with previous research indicating that factors within the tumor microenvironment, such as ascitic fluid, can impair NK cell recognition and killing abilities ^30,31^. Overcoming these inhibitory effects remains a critical challenge in improving NK cell therapies for tumors.

Interestingly, exNK-asc cells, which were exposed to suppressive conditions following expansion, retained a significant degree of cytotoxic activity (32.5%) and serial killing capacity (∼28%). Although their performance was lower than that of exNK cells, it was still markedly higher than that of pbNK and pbNK-asc cells. This suggests that while suppression following expansion diminishes NK cell function, it does not completely negate the benefits gained through the expansion process ^32^. These findings highlight the potential for combining NK cell expansion strategies with supportive treatments to counteract the suppressive effects of the tumor microenvironment.

The attachment behavior of NK cells to K562 target cells revealed critical differences in functionality across NK cell phenotypes. Enhanced target recognition and stable interaction capacity was found correlated with cytotoxic activity by pbNK-asc cells having their first attachments with targets significantly later and displaying shorter attachment durations. Notably, cytotoxic NK cells across all groups consistently demonstrated faster attachment and longer durations compared to non-cytotoxic NK cells, emphasizing the importance of stable, early attachment for effective target cell killing.

NK cell migration analysis demonstrated that motility is linked to target cell presence. Interestingly, NK cells in the presence of K562 target cells (NK+K562) migrated faster than NK cells alone (NK-K562) until attachment occurred, after which their movement slowed significantly. Since expanded NK cell groups made attachments so early, measuring their idle time velocity would be challenging. This is why they were not included in the velocity data. Moreover, NK cell migration differences might be linked to cytotoxic functionality with cytotoxic pbNK cells exhibiting higher idle time velocities compared to non-cytotoxic ones.

## 4. Conclusions / Key Points

- A droplet-based microfluidic platform was successfully employed to observe and analyze NK cell-mediated cytotoxicity at the single-cell level, and it revealed NK cell-mediated cytotoxic events and the functional diversity among NK cell phenotypes (pbNK, pbNK-asc, exNK, exNK-asc).
- The platform enabled real-time monitoring of critical events such as NK cell attachment, killing and serial killing, target cell killing times, and NK cell migration dynamics.
- exNK demonstrated superior cytotoxicity, with highest target cell killing and serial killing efficiency, and fastest killing times highlighting their potential for developing NK cell-based immunotherapies.
- pbNK-asc exhibited delayed first attachment times, shortest attachment duration, reduced killing efficiency, no serial killing activity. This reflects impaired target recognition and killing potential and, emphasizes the need for strategies to eliminate the suppressive effects of the tumor microenvironment.
- exNK-asc cells exhibited higher cytotoxicity compared to pbNK cells in all parameters. This indicates that the suppressive factors introduced after expansion do not entirely eliminate the advantages achieved through the expansion process.
- Microfluidic droplet technology is a promising tool for screening cytotoxic immune cell populations and has the potential for improving NK cell-based cancer therapies.

## 5. Methods

### 5.1 Cell Culture

NucLight Red (derived from Entacmaea quadricolor) K562 cells (created in Ashkar Lab) were maintained in RPMI-1640 (ATCC) medium supplemented with 10% FBS (ATCC), 1% Penicillin-Streptomycin (Sigma-Aldrich) and 0.1% Plasmocin^TM^ (Invivogen). Cells were grown at 37 °C and 5% CO2 in a humidified atmosphere. Cells were routinely passaged every 2 days and harvested at a 1×10^6^ viable cells/mL density.

NK cells were isolated from fresh blood of three female donors using a custom-made NK cell negative selection kit (Miltenyi Biotec, Germany) according to the manufacturer’s protocol ^33^. Purity of NK cells were confirmed using staining with antibodies against hCD45, hCD56, and hCD3 followed by flowcytometry. The isolated NK cells were cultured in RPMI-1640 medium containing 10% FBS, 1% streptomycin, 0.1% antimycotic, and 50 ng/mL IL-2. Freshly isolated pbNK cells were used immediately for assays, while portions of pbNK cells were reserved for subsequent treatments. To generate pbNK-asc cells, pbNK cells were cultured in ascites fluid collected under sterile conditions from patients with advanced ovarian cancer, as previously described ^30,31^ for three days. Ascites fluid was centrifuged at 300G for 10 minutes to remove debris prior to use. For expanded NK cells (exNK), pbNK cells were co-cultured with irradiated K562 cells expressing membrane-bound interleukin (IL-21 and recombinant human (rh) IL-2 for three weeks following established protocols ^29^. exNK-asc cells were then prepared by culturing these expanded NK cells in ascites fluid for an additional three days. All blood draws, NK cell isolation, culturing, and preparation were performed and kindly provided by Ashkar Lab (McMaster University, Hamilton, ON, Canada). All experiments were conducted with biohazard safety approval (BUP#146) and with human ethics approval from the Research Ethics Board (REB) at McMaster University (15552). Human samples were collected with written informed consent.

### 5.2 NK Cell-Mediated Cytotoxicity Assay with Droplet-based Microfluidic

Fluigent Flow EZ™ platform with appropriate tubing and connectors was employed according to the manufacturer’s instructions to encapsulate cells using droplet microchip. The encapsulation process was performed using Fluidics719 droplet generation and storage chips, which facilitate the capturing and incubation of the cells. As the continuous flow medium, dSurf, provided at 2% in 3M™ Novec™ 7500 fluorinated oil, was used. The dispersed flow consisted of 2×1^6^ cells/mL mKate2 K562 and NK cells at a concentration of 1×1^6^ cells/mL. Cells were encapsulated within droplets at E:T ratios ranging from 1:1 to 1:4. Once encapsulated, the cells were incubated at 37°C and 5% CO^2^ in a humidified atmosphere. Nikon Eclipse Ti2-A inverted microscope was used to monitor and capture images of cell interactions within the droplets. The ND acquisition software was utilized for continuous observation of cells via time-lapse imaging. Images of these locations were obtained every 15 minutes for a total period of 10 hours.

### 5.3 Analysis

NK cell-mediated cytotoxicity analysis was conducted manually using image records in ND formats. The numbers of cancer and NK cells in each droplet were recorded on Excel, along with the instances of “attachment”, “attachment but no killing” and “attachment and killing” numbers as well as killing times for each cancer cell. Attachments, killing and serial killing percentages, and killing times were calculated for each type of NK cell and E:T ratio. NK cell velocity was measured via Fiji, manual tracking plugin.

### 5.4 Statistical Analysis

Error bars on graphs represent the standard deviation. The number of samples for each experiment is specified in figure captions, and the minimum sample size used in this study is three. Graphs were analyzed by a one-way or two-way ANOVA followed by Tukey’s multiple comparison test. GraphPad Prism version 10 software (San Diego, CA, USA) was used for statistical analysis and graphs. The routines were written in Python (version 3) programming language to generate Figure 5c ^34^.

## Supporting information

Description of Additional Supplementary Files

## Conflict of Interest

The authors declare no conflict of interest.

## Acknowledgements

This research was undertaken thanks to the Canada Research Chairs Program awards to T.F. Didar. T.F. Didar is funded by the Natural Sciences and Engineering Research Council of Canada (NSERC) through the Canada Discovery Grant and the Ontario Early Researcher Award. A. A. Ashkar is funded by the Canadian Institutes of Health Research (CIHR).

## Notes

### Competing Interest Statement

The authors have declared no competing interest.

