## Supplementary Information for "Single-Cell Analysis of NK Cell Cytotoxicity in Cancer Therapy Using Microfluidic Droplets"

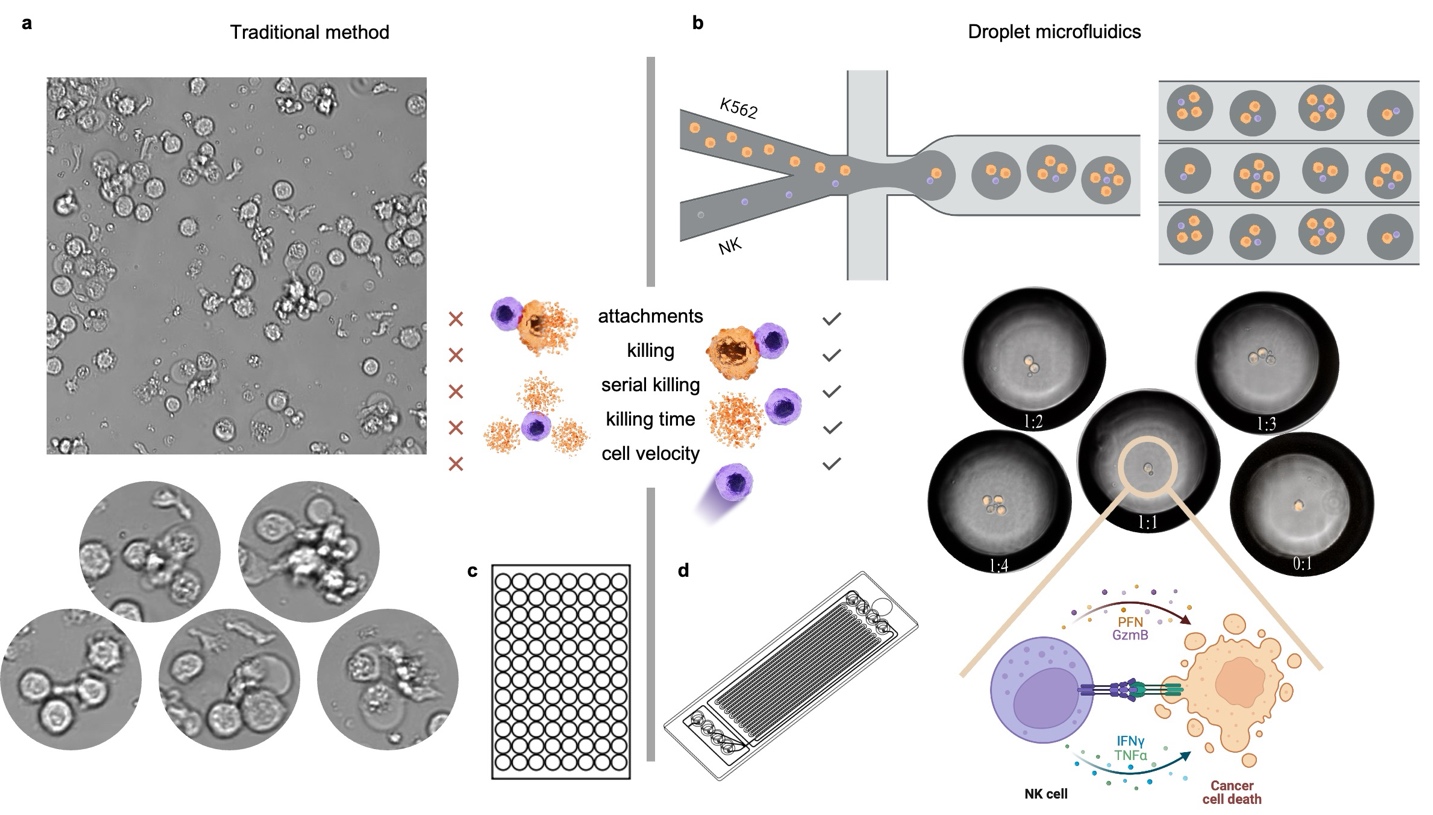

**Supplementary Figure 1: Comparative Analysis of NK Cell Cytotoxicity Assessment Methods.** (a) The traditional method of using a bulk culture of effector and target cells with uncontrolled interactions and outcomes (b) Schematic of the droplet microfluidic method and corresponding droplet images illustrating the encapsulation and precise control of E:T cell ratios, enabling targeted observations of cytotoxic activity, serial killing, and secretions (c) Image of a well plate used in conventional bulk cytotoxicity assays (d) Image of Microfluidic ChipShop's Fluidic 719 Chip used for droplet generation and storage.

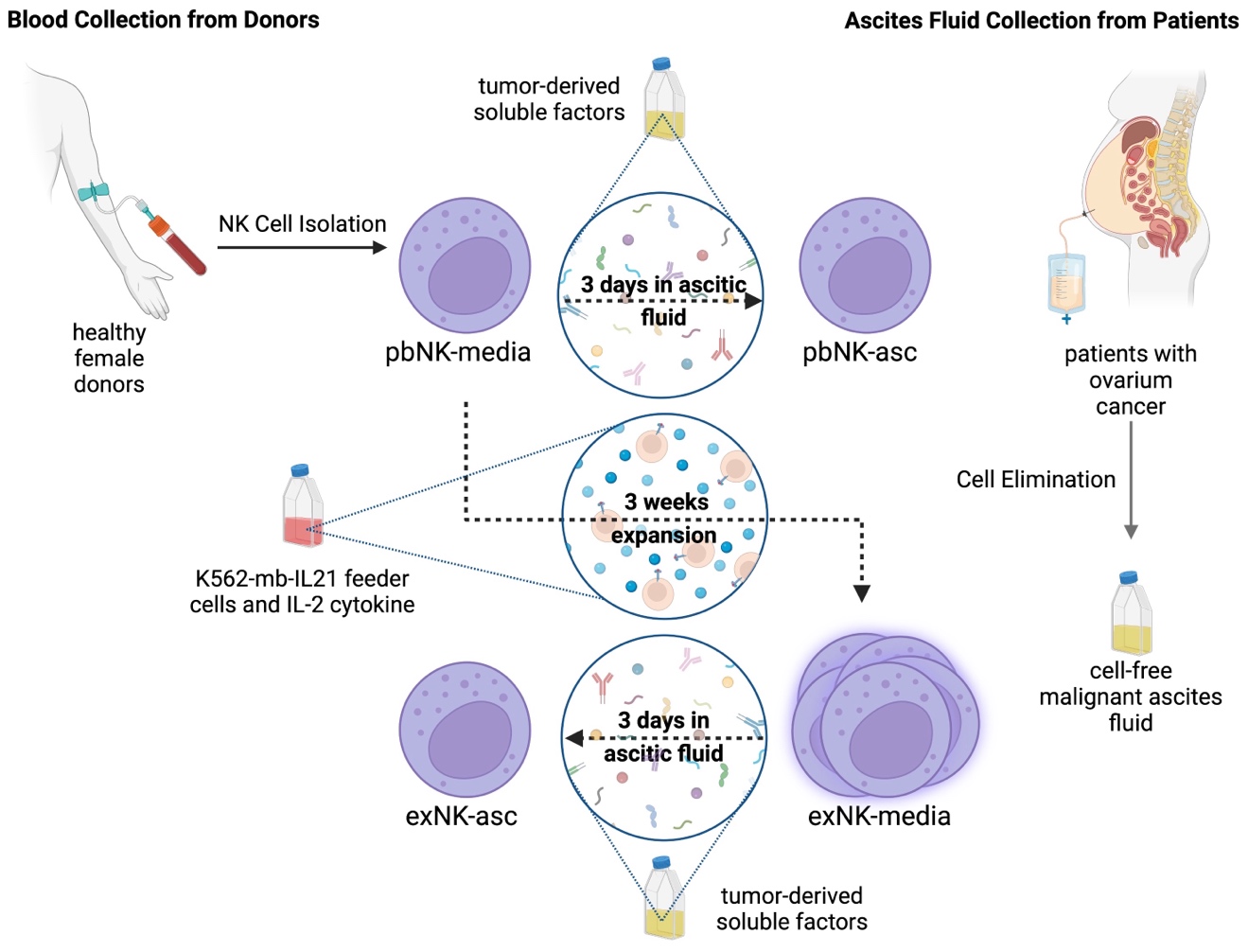

**Supplementary Figure 2: Illustration of the workflow for NK cell preparation.** NK cells are first isolated from the blood of healthy female donors. Cell-free malignant ascites fluid is collected from patients with ovarian cancer, representing the tumor-derived soluble factors that simulate the tumor microenvironment. Primary NK cells (pbNK-media) are cultured in ascites fluid for 3 days (pbNK-asc). In parallel, NK cells undergo a 3-week expansion using K562-mb-IL21 feeder cells and IL-2 cytokine to generate expanded NK cells (exNK-media). Expanded NK cells (exNK-asc) are also exposed to the ascites fluid for 3 days.

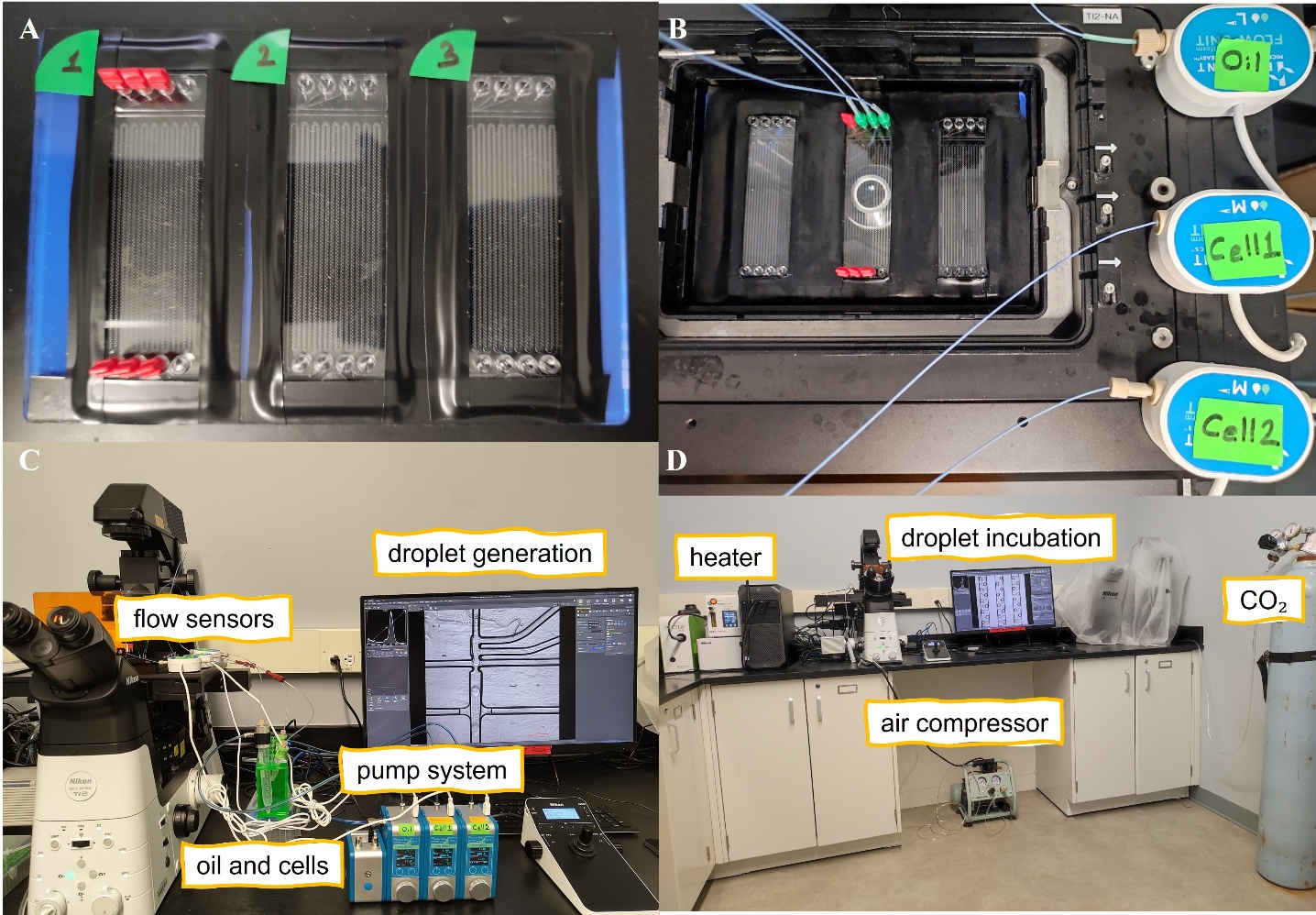

**Supplementary Figure 3: Droplet Generation and Capturing with Fluidic 719 Chip**. (a) Three Fluidics719 droplet‑generation/storage chips loaded in parallel for simultaneous processing. (b) A single chip mounted in the Fluigent flow sensors, with inlet tubing for oil and cell suspensions. (c) Droplet‑generation workstation featuring a Nikon Eclipse Ti2‑E inverted microscope, Fluigent Flow EZ pump system, and inline flow sensors, with reservoirs for dSurf oil and cell suspensions. (d) Droplet‑incubation bench showing the heated microscope enclosure and CO₂ cylinder used to maintain 37 °C and 5% CO₂ during time‑lapse imaging.

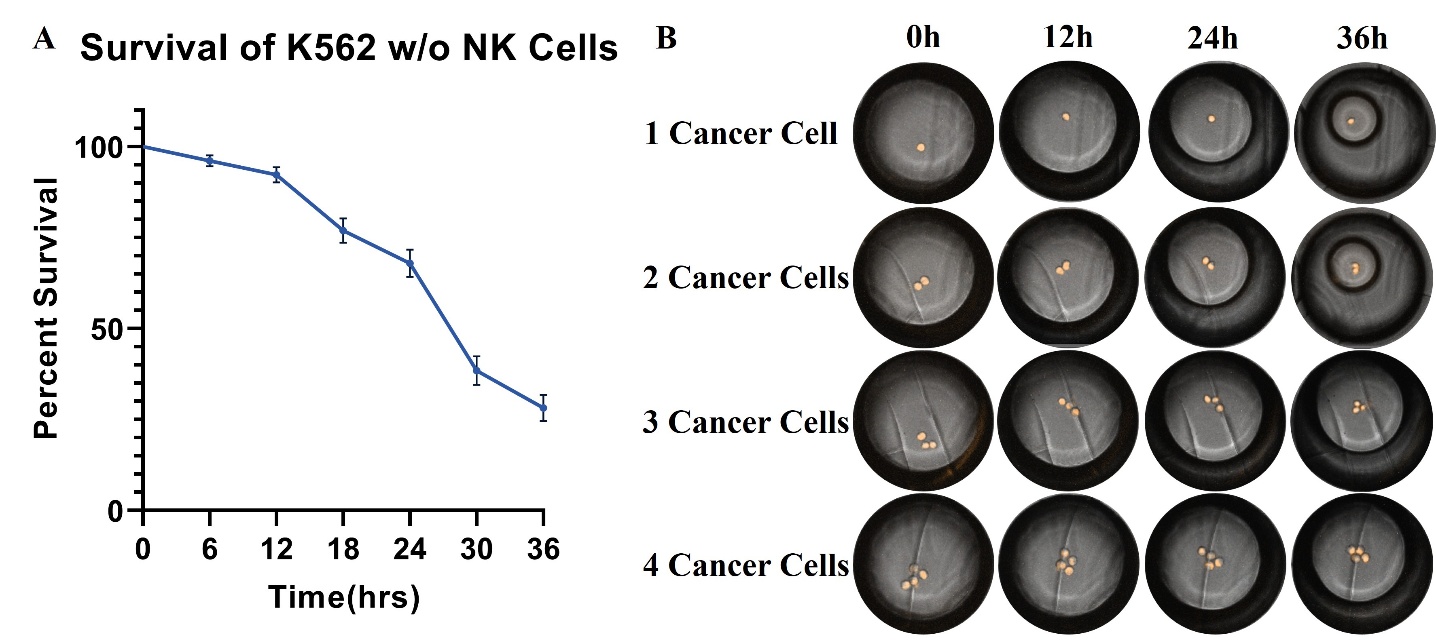

**Supplementary Figure 4: Survival of K562 Cells Without NK Cells.** (a) Graph showing the survival of K562 cells over a 36-hour incubation period in the absence of NK cells. (b) Sequential images of droplets containing different numbers of K562 cancer cells taken at 0, 12, 24, and 36 hours of incubation. These images illustrate the survivability of the cancer cells without NK cells over time within the droplets.

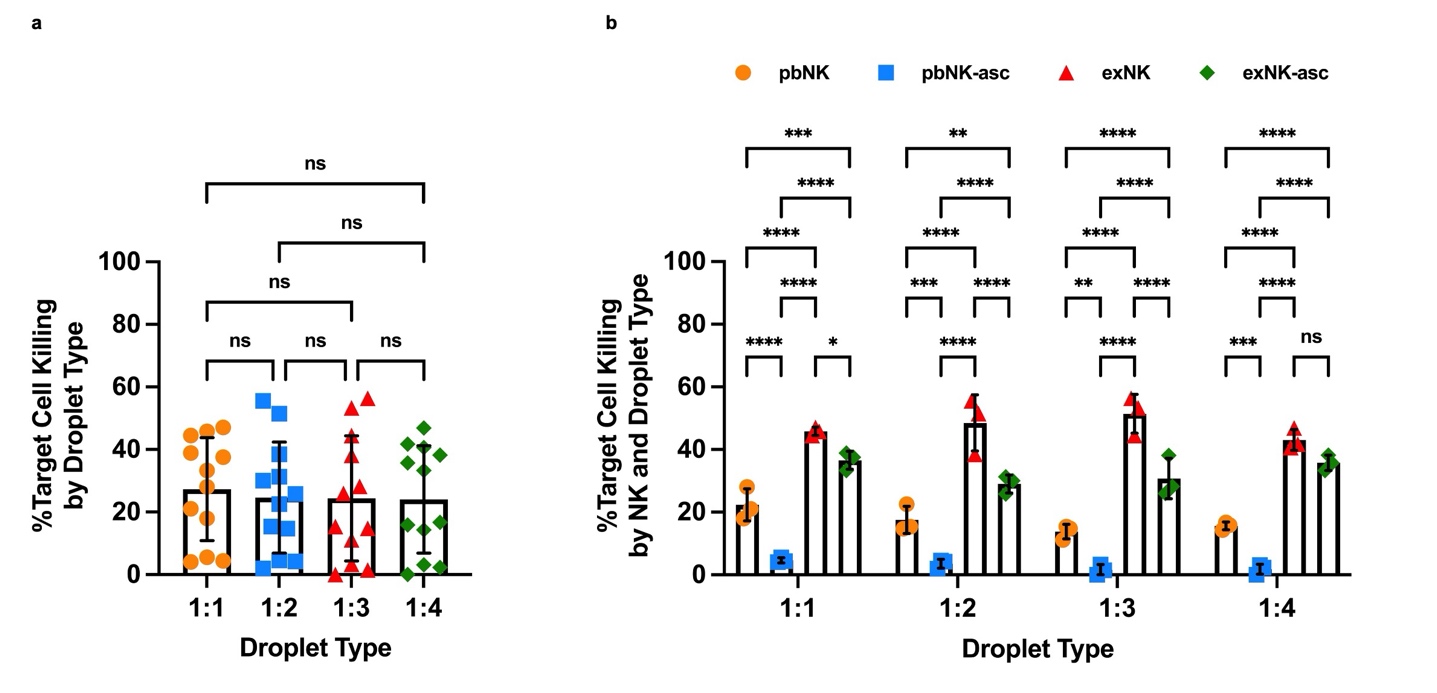

**Supplementary Figure 5: Comparison of Target Cell Killing Across Droplet Types and NK Cell Groups.** (a) Percentage of target cell killing across different droplet types (E:T ratios ranging from 1:1 to 1:4) regardless of NK cell type. (b) Percentage of target cell killing by NK cell groups (pbNK, pbNK-asc, exNK, and exNK-asc) across varying droplet types. Data represent three independent biological replicates (n=3) for each condition. Asterisks represent “ns” (p > 0.05), * (p ≤ 0.05), ** (p ≤ 0.01), *** (p ≤ 0.001), and **** (p ≤ 0.0001) from one-way or two-way ANOVA with Tukey’s multiple comparisons test. All error bars represent the standard deviation.

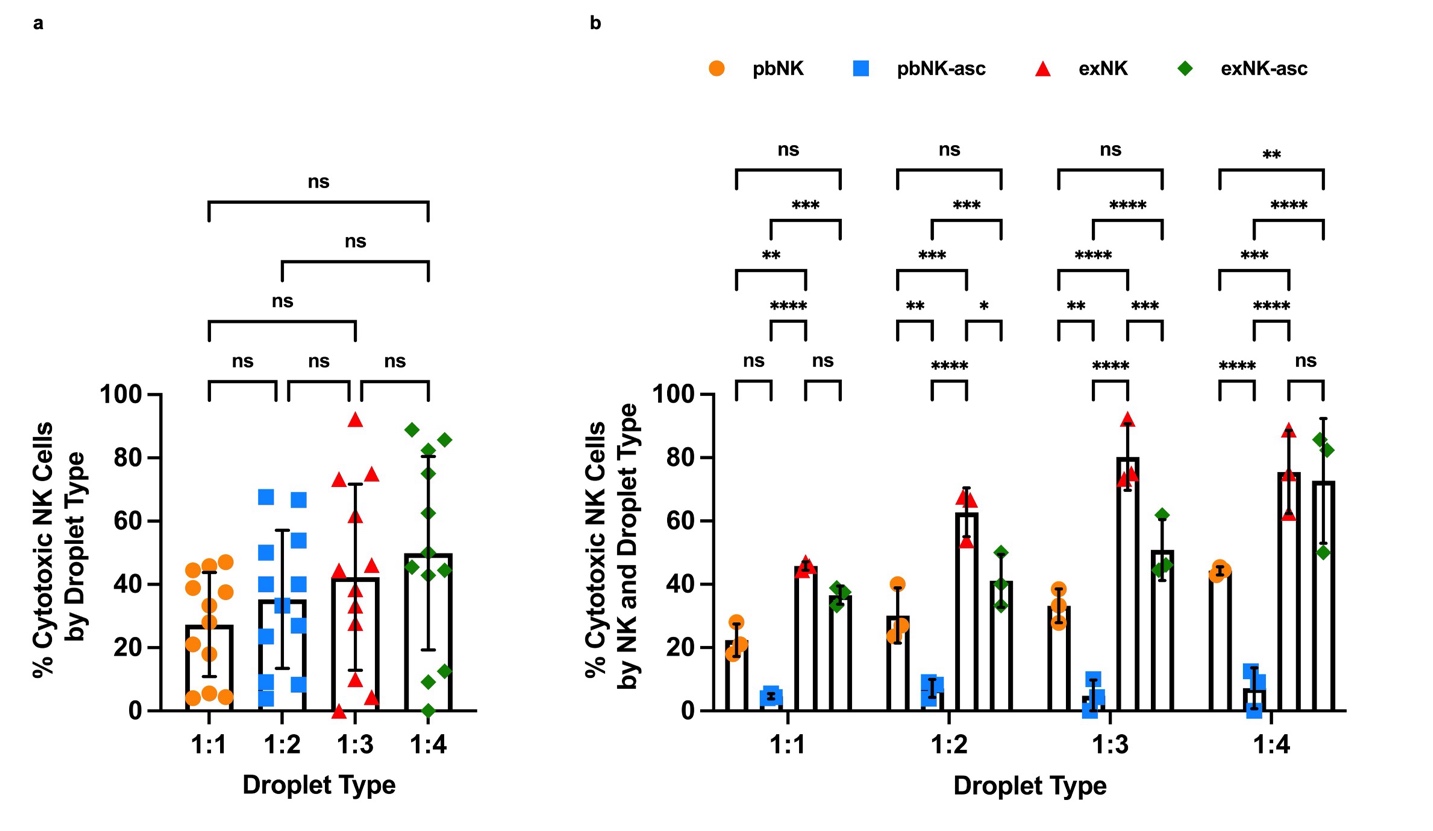

**Supplementary Figure 6: Percentage of Cytotoxic NK Cells Across Droplet Types and NK Cell Groups.** (a) Percentage of cytotoxic NK cells across different droplet types (E:T ratios ranging from 1:1 to 1:4) regardless of NK cell group. (b) Percentage of cytotoxic NK cells by NK cell groups (pbNK, pbNK-asc, exNK, and exNK-asc) across varying droplet types. Data represent three independent biological replicates (n=3) for each condition. Asterisks represent “ns” (p > 0.05), * (p ≤ 0.05), ** (p ≤ 0.01), *** (p ≤ 0.001), and **** (p ≤ 0.0001) from one-way or two-way ANOVA with Tukey’s multiple comparisons test. All error bars represent the standard deviation.

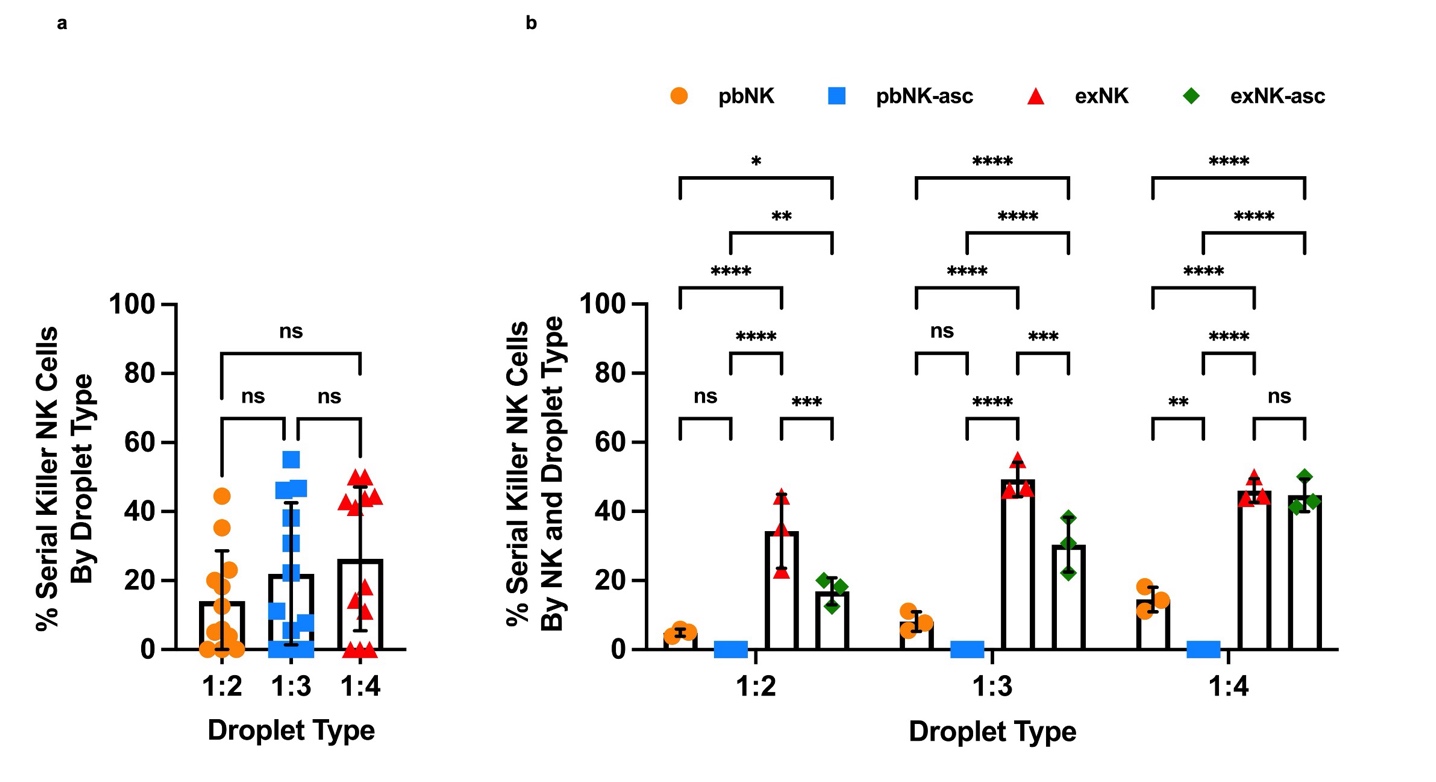

**Supplementary Figure 7: Percentage of Serial Killer NK Cells Across Droplet Types and NK Cell Groups.** (a) Percentage of serial killer NK cells across different droplet types (E:T ratios 1:2, 1:3, and 1:4) regardless of NK cell group. (b) Percentage of serial killer NK cells by NK cell groups (pbNK, pbNK-asc, exNK, and exNK-asc) across varying droplet types. Statistical significance was determined using one-way ANOVA with Tukey's multiple comparisons test. Asterisks represent * (p ≤ 0.05), ** (p ≤ 0.01), *** (p ≤ 0.001), and **** (p ≤ 0.0001), while ns indicates non-significant comparisons. Error bars represent the standard deviation.

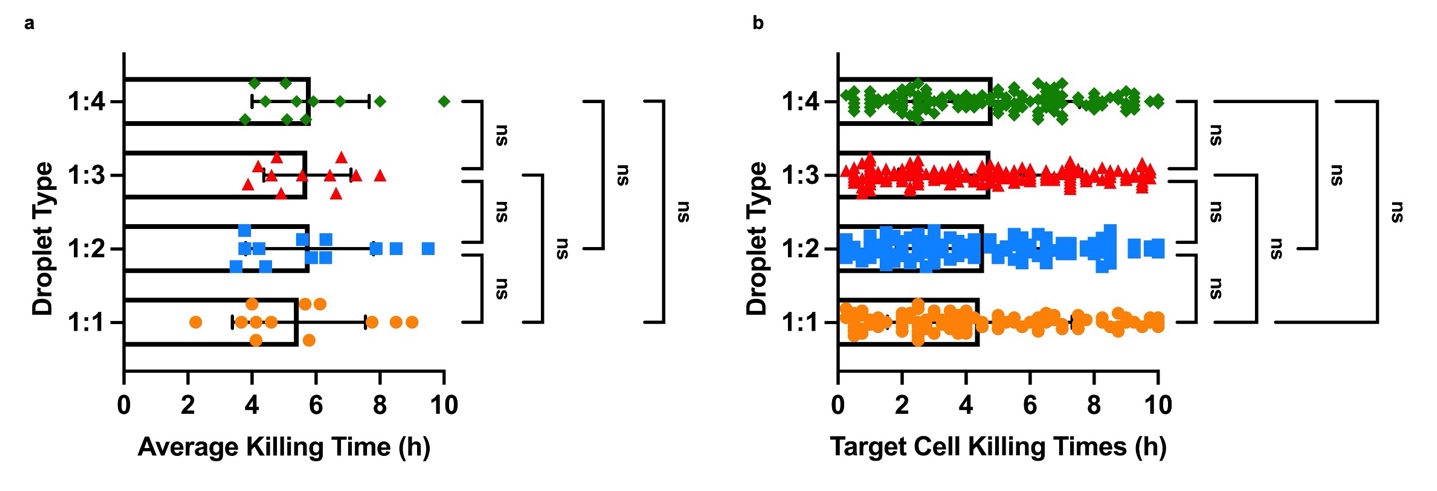

**Supplementary Figure 8:** **Supplementary Figure 8: Comparison of Killing Times Across Droplet Types.** (a) Average target cell killing times across different droplet types (E:T ratios ranging from 1:1 to 1:4) for all NK cell groups combined. (b) Distribution of individual target cell killing times across varying droplet types (E:T ratios). Data represent three independent biological replicates (n=3) for each condition. Asterisks represent “ns” (p > 0.05), * (p ≤ 0.05), ** (p ≤ 0.01), *** (p ≤ 0.001), and **** (p ≤ 0.0001) from one-way or two-way ANOVA with Tukey’s multiple comparisons test. All error bars represent the standard deviation.

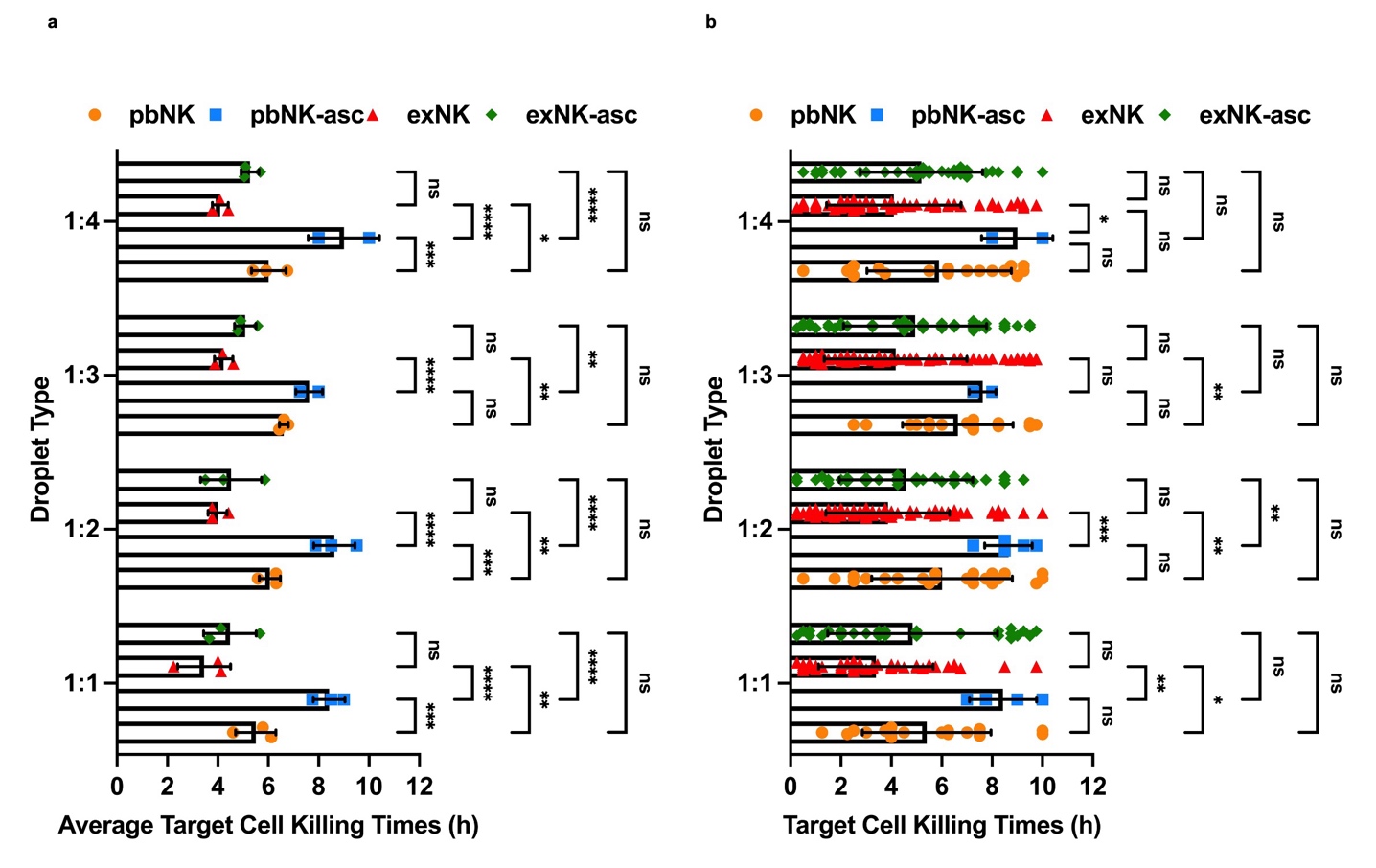

**Supplementary Figure 9: Supplementary Figure 9: Comparison of Target Cell Killing Times Across Droplet Types and NK Cell Groups.** (a) Average target cell killing times for different NK cell groups (pbNK, pbNK-asc, exNK, and exNK-asc) across varying droplet types (E:T ratios ranging from 1:1 to 1:4). (b) Distribution of individual target cell killing times for each NK cell group across droplet types. Data represent three independent biological replicates (n=3) for each condition. Asterisks represent “ns” (p > 0.05), * (p ≤ 0.05), ** (p ≤ 0.01), *** (p ≤ 0.001), and **** (p ≤ 0.0001) from one-way or two-way ANOVA with Tukey’s multiple comparisons test. All error bars represent the standard deviation.

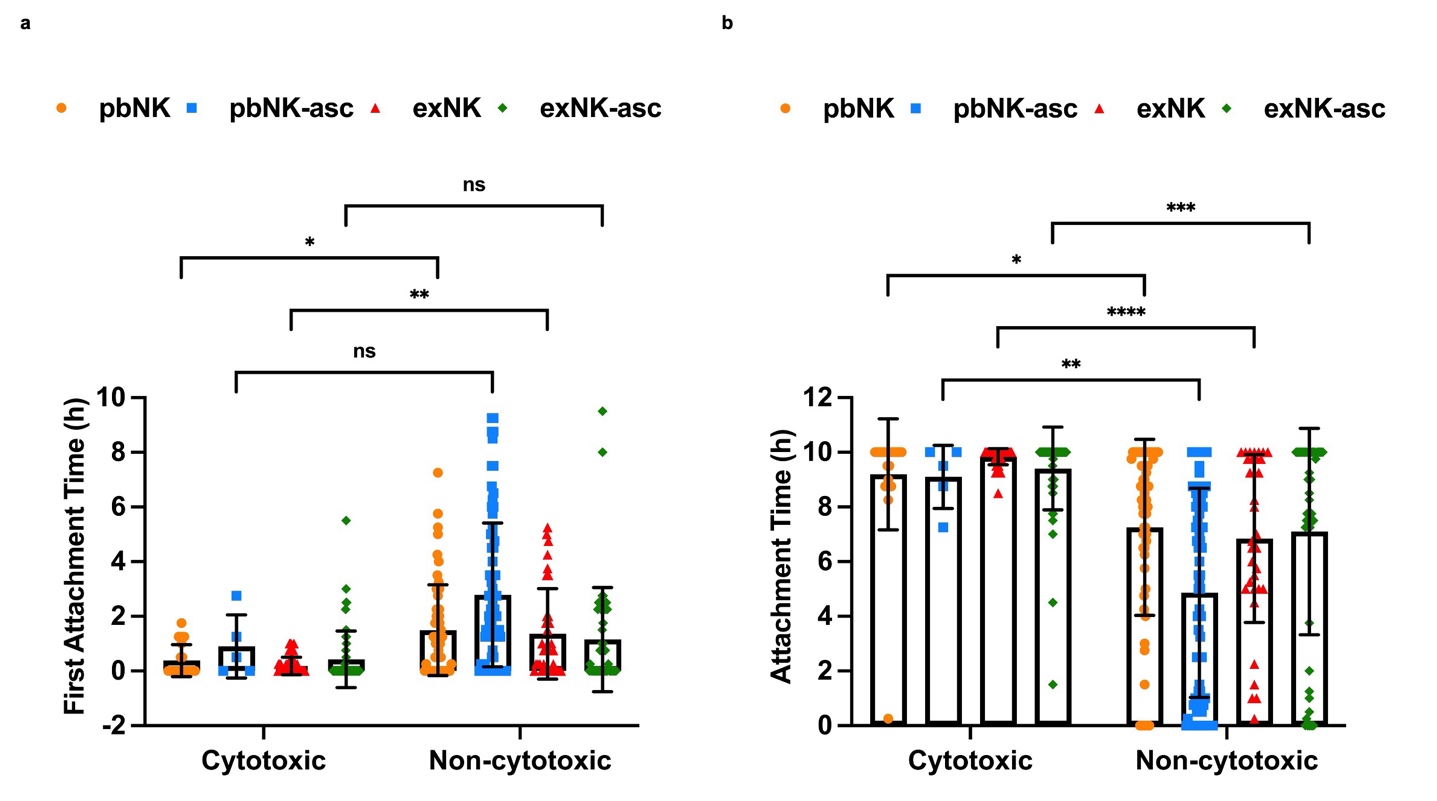

**Supplementary Figure 10: Supplementary Figure 10: Comparison of NK Cell Attachment Times Across NK Cell Groups.** (a) First attachment times of cytotoxic and non-cytotoxic NK cells (pbNK, pbNK-asc, exNK, and exNK-asc) with K562 target cells. (b) Total attachment times of cytotoxic and non-cytotoxic NK cells across NK cell groups. Data represent three independent biological replicates (n=3) for each condition. Asterisks represent “ns” (p > 0.05), * (p ≤ 0.05), ** (p ≤ 0.01), *** (p ≤ 0.001), and **** (p ≤ 0.0001) from one-way or two-way ANOVA with Tukey’s multiple comparisons test. All error bars represent the standard deviation.

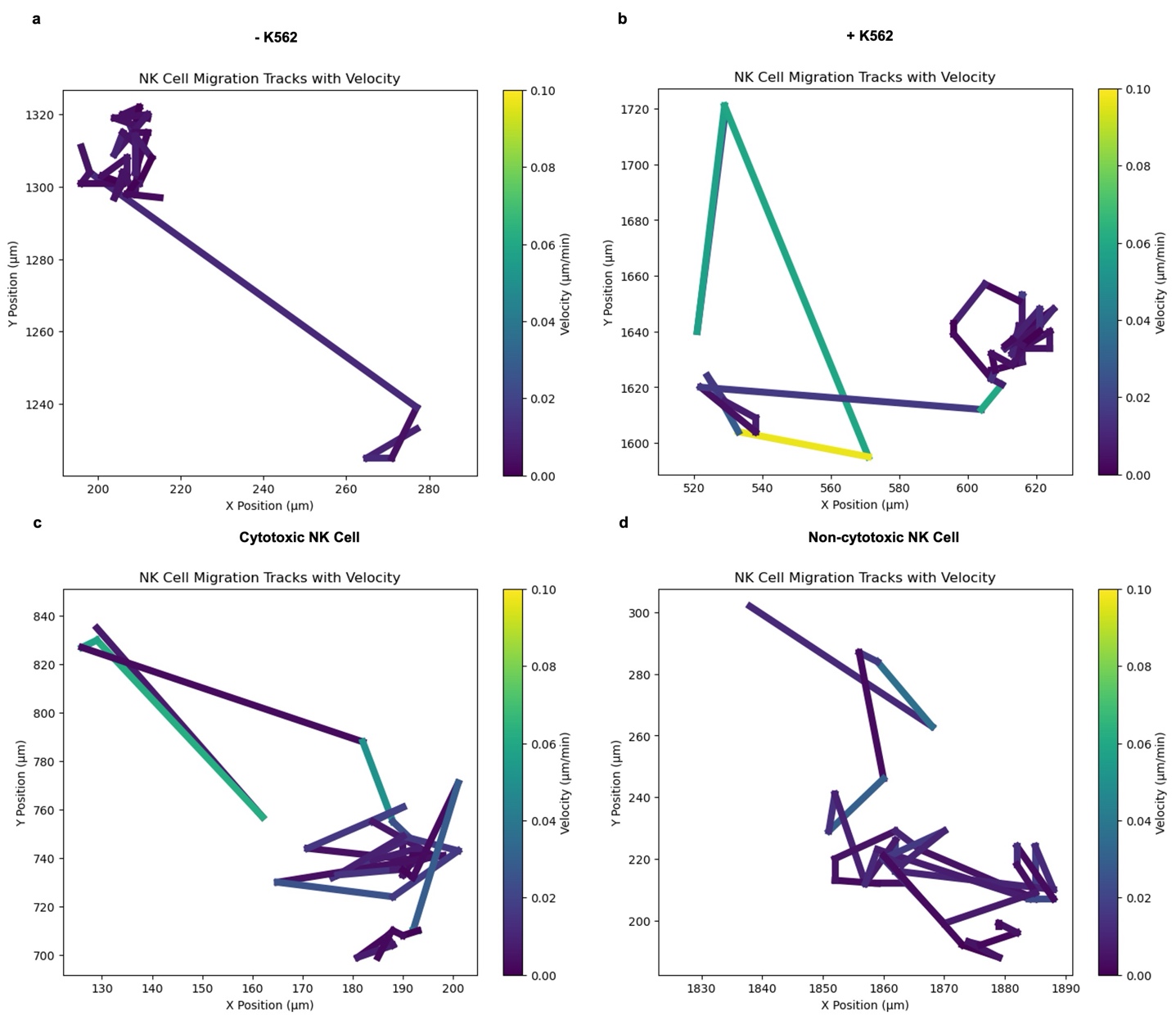

**Supplementary Figure 11:** **NK Cell Migration Tracks and Instantaneous Velocity.** (a) Representative migration tracks of NK cells alone (-K562) and (b) in the presence of K562 target cells (+K562). (c) Migration tracks of cytotoxic NK cells showing higher motility before attachment. (d) Migration tracks of non-cytotoxic NK cells, indicating reduced movement compared to cytotoxic NK cells. The colormap represents the instantaneous velocity of NK cells during migration (µm/min).

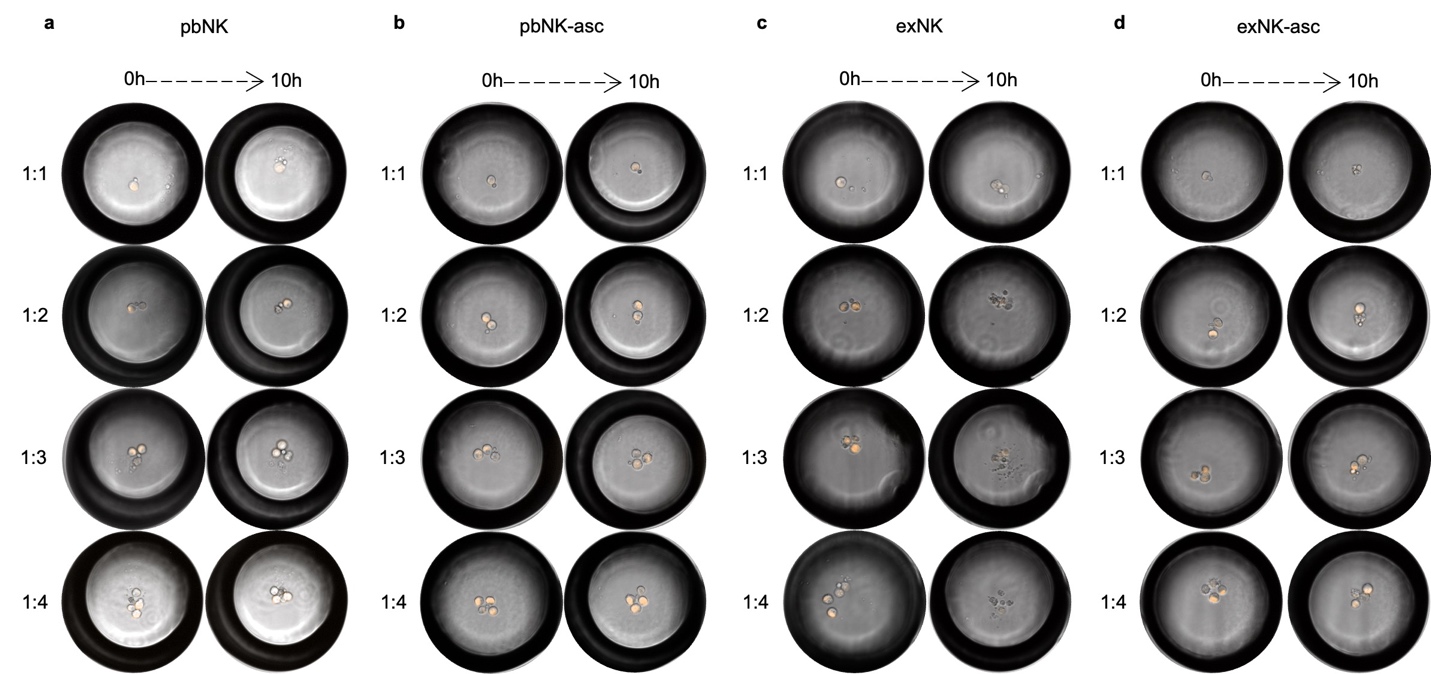

**Supplementary Figure 12:** Time-Lapse Analysis of NK Cell-Mediated Cytotoxicity in Droplet Microfluidics. Illustration of the interaction dynamics of (a) pbNK, (b) pbNK-asc, (c) exNK and (d) exNK-asc cells with K562 over 10 hours at various E:T ratios, highlighting differences in cytotoxic engagement and the impact of cell conditioning on NK cell activity.
