## Supplementary material for "Single-Cell Analysis of NK Cell Cytotoxicity in Cancer Therapy Using Microfluidic Droplets": Description of Additional Supplementary Files

File Name: Supplementary Movie 1

Description: Droplet generation and capturing with Fluidic 719 chip. Droplets are generated in a flow-focusing channel to encapsulate NK and K562 cells with continuous oil flow. Generated droplets flow into droplet capturing area for long-term incubation. The droplets are directed into the traps with capillary pressure via geometric constrictions.

File Name: Supplementary Movie 2

Description: NK and K562 cells are encapsulated at a 1:4 effector-to-target ratio, illustrating different killing outcomes over 10 hours. Top-left: peripheral blood NK cells (pbNK) kill 1 of 4 target cells. Top-right: TME-conditioned pbNK (pbNK-asc) kill none (0 of 4). Bottom-left: expanded NK cells (exNK) kill all 4 targets. Bottom-right: TME-conditioned exNK (exNK-asc) kill 2 of 4 targets.

File Name: Supplementary Movie 3

Description: NK and K562 target cells are encapsulated to compare NK cell velocities when they are alone and when they are with a target as well as to compare cytotoxic and non-cytotoxic NK cell velocities. Top-left: NK cell alone (NK(-K562)). Top-right: NK cell with a K562 target (NK(+K562)). Bottom-left: a cytotoxic NK cell killing K562. Bottom-right: a non-cytotoxic NK cell that does not kill its target.
